# Mitochondrial impairment activates the Wallerian pathway through depletion of NMNAT2 leading to SARM1-dependent axon degeneration

**DOI:** 10.1101/683342

**Authors:** Andrea Loreto, Ciaran S. Hill, Victoria L. Hewitt, Giuseppe Orsomando, Carlo Angeletti, Jonathan Gilley, Cristiano Lucci, Alvaro Sanchez-Martinez, Alexander J. Whitworth, Laura Conforti, Federico Dajas-Bailador, Michael P. Coleman

## Abstract

Wallerian degeneration of physically injured axons involves a well-defined molecular pathway linking loss of axonal survival factor NMNAT2 to activation of pro-degenerative protein SARM1. Manipulating the pathway through these proteins led to the identification of non-axotomy insults causing axon degeneration by a Wallerian-like mechanism, including several involving mitochondrial impairment. Mitochondrial dysfunction is heavily implicated in Parkinson’s disease, Charcot-Marie-Tooth disease, hereditary spastic paraplegia and other axonal disorders. However, whether and how mitochondrial impairment activates Wallerian degeneration has remained unclear. Here, we show that disruption of mitochondrial membrane potential leads to axonal NMNAT2 depletion in mouse sympathetic neurons, increasing the substrate-to-product ratio (NMN/NAD) of this NAD-synthesising enzyme, a metabolic fingerprint of Wallerian degeneration. The mechanism appears to involve both impaired NMNAT2 synthesis and reduced axonal transport. Expression of WLD^S^ and *Sarm1* deletion both protect axons after mitochondrial uncoupling. Blocking the pathway also confers neuroprotection and increases the lifespan of flies with *Pink1* loss-of-function mutation, which causes severe mitochondrial defects. These data indicate that mitochondrial impairment replicates all the major steps of Wallerian degeneration, placing it upstream of NMNAT2 loss, with the potential to contribute to axon pathology in mitochondrial disorders.

## INTRODUCTION

Studies of axon degeneration following axotomy (Wallerian degeneration) and of the axon-protective protein WLD^S^ have led to the discovery of critical endogenous regulators of the mechanisms resulting in axon degeneration (Conforti et al., 2014; Gerdts et al., 2016). The current model predicts that the pathway regulating Wallerian degeneration (Wallerian pathway) is activated by the loss in the axon of the labile nicotinamide mononucleotide adenylyl-transferase 2 (NMNAT2), a nicotinamide adenine dinucleotide (NAD)-synthesising enzyme. Axonal NMNAT2 levels decline within a few hours when its transport and/or synthesis are impaired (Gilley and Coleman, 2010). Downstream of NMNAT2 depletion, the pro-degenerative protein sterile alpha and TIR motif-containing protein 1 (SARM1) executes the degeneration program (Gerdts et al., 2015; Gilley et al., 2015; Loreto et al., 2015; Osterloh et al., 2012). To date, expression of WLD^S^/NMNATs (which substitute for endogenous NMNAT2 loss) and SARM1 depletion are the most effective means to block the Wallerian pathway and preserve axons in mammals. There is still debate about how NMNAT2 loss leads to SARM1 activation but the rise in its substrate, NMN, appears to be important (Cohen, 2017; Di Stefano et al., 2015, 2017; Loreto et al., 2015; Zhao et al., 2019) as well as the fall in its product, NAD (Essuman et al., 2017; Gerdts et al., 2015; Sasaki et al., 2016).

Most studies on the Wallerian pathway have used a physical injury model, but there is clear evidence that related degenerative mechanisms can be activated by many non-injury stresses (Conforti et al., 2014). However, the vast majority of non-axotomy models were performed when a more comprehensive understanding of the Wallerian pathway was lacking and were mostly identified as being Wallerian-like by targeting just a single step in the pathway (either by expressing of WLD^S^/NMNATs or, more recently, by *Sarm1* deletion). Both WLD^s^ and SARM1 have the potential to influence other cellular mechanisms, such as nuclear NAD synthesis and innate immunity, respectively, so involvement of the Wallerian pathway is best supported by multiple lines of evidence.

The link between mitochondria and the Wallerian pathway is particularly intriguing. Mitochondrial dysfunction is a common theme in a wide group of neurodegenerative disorders in which axon degeneration is central, including Parkinson’s disease (PD), Charcot-Marie-Tooth disease, hereditary spastic paraplegia and Friedrich’s ataxia (Court and Coleman, 2012). We and others have previously shown that mitochondria contribute to the later stages of Wallerian degeneration, where the axotomy itself activates the Wallerian pathway (Barrientos et al., 2011; Loreto et al., 2015). However, mitochondrial depolarisation, caused by the mitochondrial uncoupler Carbonyl cyanide m-chlorophenyl hydrazone (CCCP), also leads to degeneration of uninjured axons (Loreto et al., 2015), which is rescued by *Sarm1* deletion (Summers et al., 2014). Additional studies, both *in vitro* and *in vivo*, link the Wallerian pathway to mitochondrial impairment. *Wld*^*S*^ mice are protected against nigrostriatal axon degeneration after intraperitoneal administration of the mitochondrial complex-1 inhibitor 1-methyl-4-phenyl-1,2,3,6-tetrahydropyridine (MPTP) (Hasbani and O’Malley, 2006). WLD^S^ also preserves neurites and promotes neuronal survival in primary dopaminergic neurons treated with MPP^+^ (the active metabolite of MPTP) (Antenor-Dorsey and O’Malley, 2012). Finally, NMNATs overexpression and *Sarm1* deletion in sensory neurons delay axon degeneration caused by rotenone, another mitochondrial complex-1 inhibitor, in sensory neurons (Press and Milbrandt, 2008; Summers et al., 2014). Thus, we hypothesised that mitochondrial impairment can also act as an upstream cause, equivalent to physical injury, in initiating the Wallerian pathway.

Here we combine multiple lines of evidence to firmly establish a role for the Wallerian pathway in axon degeneration caused by mitochondrial depolarisation in the absence of a physical injury. We also corroborate these findings using an *in vivo* genetic model of mitochondrial dysfunction, reporting a neuroprotective role of regulators of Wallerian degeneration in dopaminergic neuron loss in *Pink1* mutant flies.

## RESULTS

### Multiple regulators of the Wallerian pathway rescue axon degeneration caused by mitochondrial depolarisation

The mitochondrial uncoupler CCCP is widely used to trigger mitochondrial depolarisation and assess the effects of mitochondrial impairment on cellular viability (Ly et al., 2003). Previous work by us and others demonstrated that sympathetic and sensory primary neurons exposed to CCCP undergo disruption of mitochondrial membrane potential and axon degeneration (Loreto et al., 2015; Summers et al., 2014), providing a good experimental model to study mitochondrial dysfunction leading to axon degeneration.

A dose-response experiment in superior cervical ganglion (SCG) neurons confirmed previous observations and allowed us to determine the most appropriate concentration of CCCP to use across the study. 50µM CCCP induces full mitochondrial depolarization within minutes after its addition (Loreto et al., 2015) and a dramatic depletion of ATP levels within the first 2 hr (Fig. 1A). Importantly, it consistently promoted neurite degeneration when measured at 24 hr post-application (Fig. 1B-G). We then tested whether this degenerative process could be rescued by regulators of the Wallerian pathway. Consistent with previous studies (Summers et al., 2014), we found that *Sarm1*^*-/-*^ SCG neurites were strongly protected against CCCP toxicity (Fig. 1D, E). WLD^S^ expression was highly protective too at this concentration (Fig. 1F, G). Our findings demonstrate that the degeneration of axons following mitochondrial depolarisation can be delayed by multiple regulators of the Wallerian pathway.

**Figure 1.**
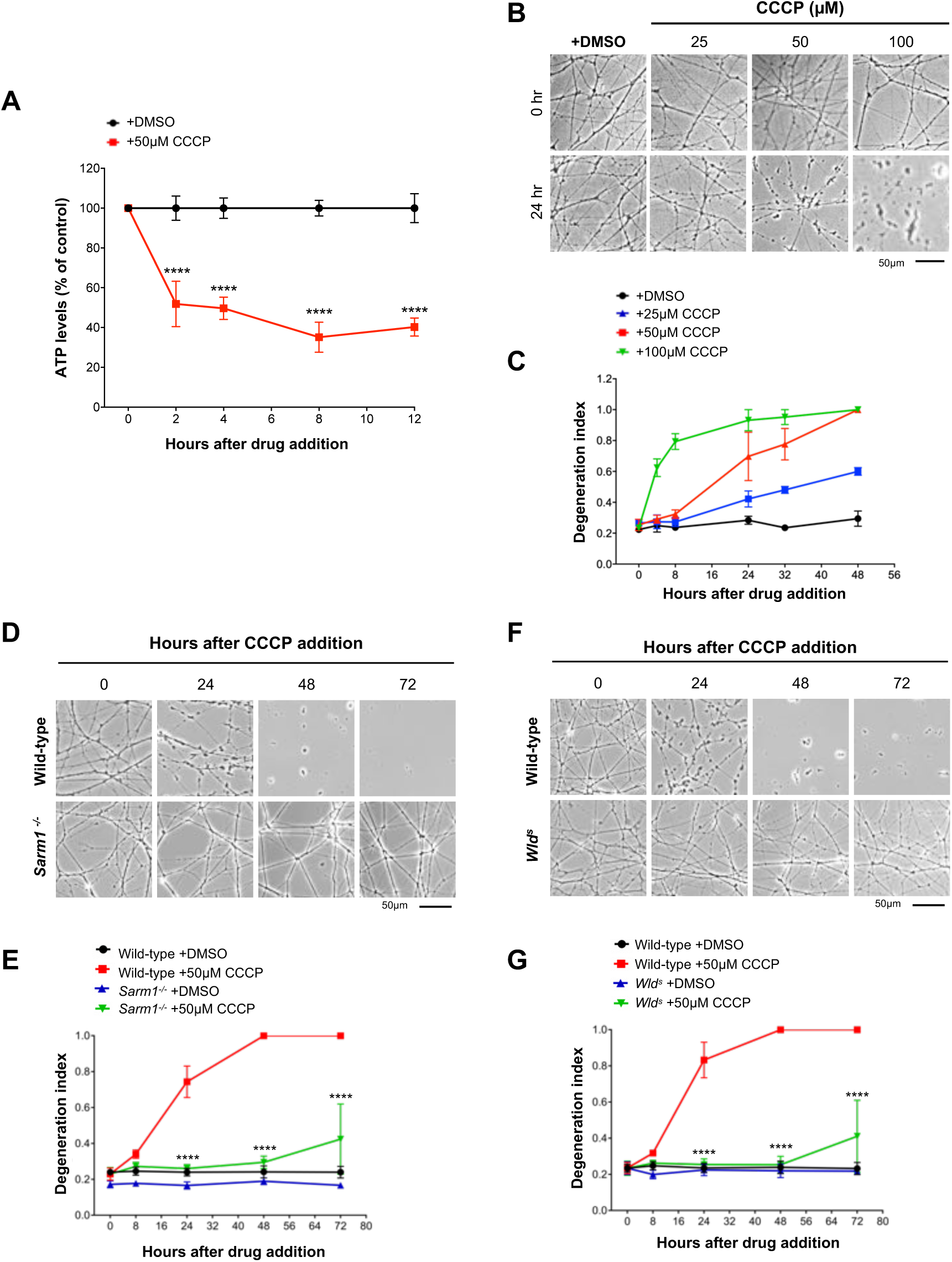
Regulators of the Wallerian pathway rescue axon degeneration caused by mitochondrial depolarisation. **(A)** ATP levels in wild-type SCG dissociated cultures after treatment with CCCP. Data are normalised to DMSO control at each time point (Mean±SEM; n=4; two-way ANOVA followed by Sidak post-hoc test; ****, p<0.0001). **(B)** Representative phase contrast images of neurites from wild-type SCG explant cultures treated with increasing concentrations of CCCP. **(C)** Quantification of the degeneration index in experiments described in (B) from 3 fields per sample in 2 independent experiments (Mean±SEM; n=2). **(D)** Representative phase contrast images of neurites from wild-type and *Sarm1*^*-/-*^ SCG explant cultures at the indicated time points after CCCP treatment. **(E)** Quantification of the degeneration index in experiments described in (D) from 3 fields per sample in 4 independent experiments (Mean±SEM; n=4; two-way ANOVA followed by Tukey post-hoc test; ****, p<0.0001. Statistical significance shown relative to +50 µM CCCP). **(F)** Representative phase contrast images of neurites from wild-type and *Wld*^*s*^ SCG explant cultures at the indicated time points after CCCP treatment. **(G)** Quantification of the degeneration index in experiments described in (F) from 3 fields per sample in 4 independent experiments (Mean±SEM; n=4; two-way ANOVA followed by Tukey post-hoc test; ****, p<0.0001. Statistical significance shown relative to +50µM CCCP).

### Mitochondrial depolarisation leads to depletion of axonal NMNAT2

NMNAT2 depletion in axons has been proposed as an initial step that triggers the activation of the Wallerian pathway (Gilley and Coleman, 2010; Gilley et al., 2015; Loreto et al., 2015; Walker et al., 2017). We therefore tested whether CCCP treatment led to NMNAT2 depletion in neurites (which were uninjured until immediately prior to harvesting separately from their cell bodies) and found that levels of this protein in neurites rapidly decline from 2 hr after CCCP addition (Fig. 2A, B). Loss of NMNAT2 occurred before any visible morphological damage to neurites (Fig. 2C), also confirmed by the absence of changes to β-actin levels (Fig. 2A). Levels of SCG10, another short-lived protein comigrating with NMNAT2 (Milde et al., 2013) and involved in Wallerian degeneration (Shin et al., 2012) and sporadic ALS (Melamed et al., 2019), declined with a similar timecourse (Fig. 2A, B).

**Figure 2.**
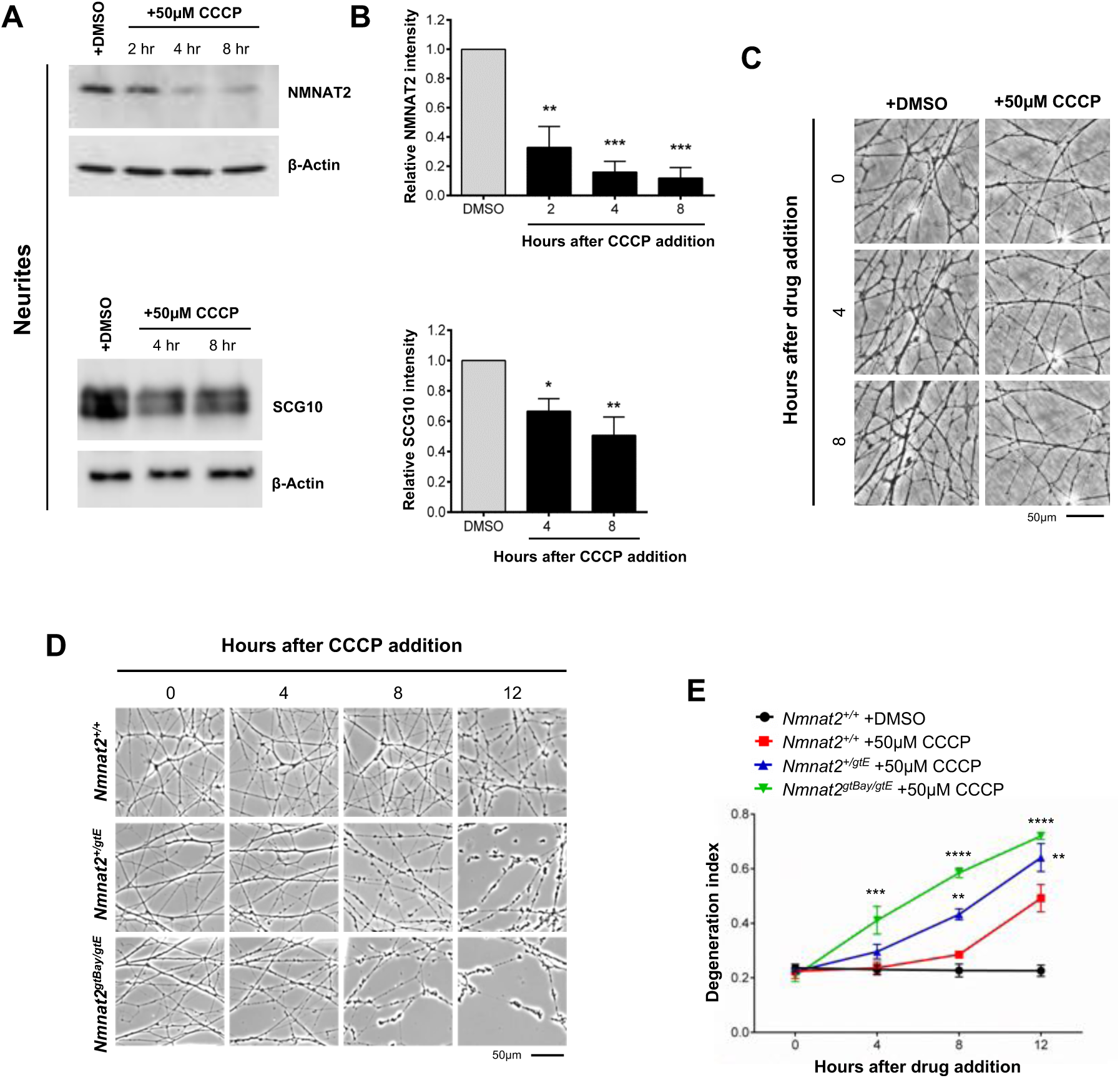
Mitochondrial depolarisation leads to depletion of axonal NMNAT2. **(A)** Representative immunoblots of wild-type SCG neurite extracts probed for NMNAT2, SCG10 and β-actin (loading control) at the indicated time points after CCCP treatment. **(B)** Quantification of normalised NMNAT2 and SCG10 levels (to β-actin) is shown, with data presented relative to DMSO control (Mean±SEM; n=3-4; one-way ANOVA followed by Bonferroni post-hoc test; ***, p<0.001; **, p<0.01; *, p<0.05). **(C)** Representative phase contrast images showing morphologically intact neurites at the time points used in (A). **(D)** Representative phase contrast images of neurites from wild-type-*Nmnat2*^*+/+*^, *Nmnat2*^*+/gtE*^ (∼60% expression), *Nmnat2*^*gtBay/gtE*^ (∼30% expression) SCG explant cultures at the indicated time points after CCCP treatment. **(E)** Quantification of the degeneration index in experiments described in (D) from 3 fields per sample in 4 independent experiments (Mean±SEM; n=4; two-way ANOVA followed by Tukey post-hoc test; ****, p<0.0001; ***, p<0.001; **, p<0.01. Statistical significance shown relative to *Nmnat2*^*+/+*^ +50µM CCCP).

We have recently reported that lowering the expression of NMNAT2 increases axonal vulnerability to several stresses (Gilley et al., 2019). To test whether lowering NMNAT2 expression impairs the ability to withstand mitochondrial impairment, SCG neurons from mice with around 60% (*Nmnat2*^*+/gtE*^) and 30% (*Nmnat2*^*gtBay/gtE*^) of wild type *Nmnat2* mRNA levels in whole brain (Gilley et al., 2019) were exposed to CCCP. We found a significant acceleration of the degeneration process compared to wild type neurons, with clear morphological damage appearing as early as 4 hr in *Nmnat2*^*gtBay/gtE*^ neurites (Fig. 2D, E).

These data suggest that mitochondrial uncoupling activates the Wallerian pathway at an early step and, together with the protection afforded by WLD^S^ (Fig. 1F, G), they indicate that axonal NMNAT levels modulate axon survival after mitochondrial depolarisation.

### NMNAT2 depletion reflects impairment of both axonal transport and synthesis

We next investigated the cause of NMNAT2 depletion after CCCP treatment. Being a labile protein with a half-life of less than an hour (Milde et al., 2013), any cellular process that impairs its replenishment in axons would lead to a rapid decrease in axonal levels. Two potential mechanisms are a deficiency in axonal transport and/or altered synthesis, both of which are ATP-dependent. The finding that NMNAT2 levels also declined in the cell body/ganglia fraction after 4-8 hr of CCCP addition (Fig. 3A, B) suggests that synthesis of the protein is impaired (although enhanced protein degradation cannot be ruled out). However, the NMNAT2 decrease in the cell body fraction was much less marked than that in neurites (Fig. 2A, B), suggesting that impaired protein synthesis is not the only mechanism contributing to the depletion in the neurites. SCG10 levels in the cell body fraction, instead, did not vary significantly (Fig. 3A, B).

**Figure 3.**
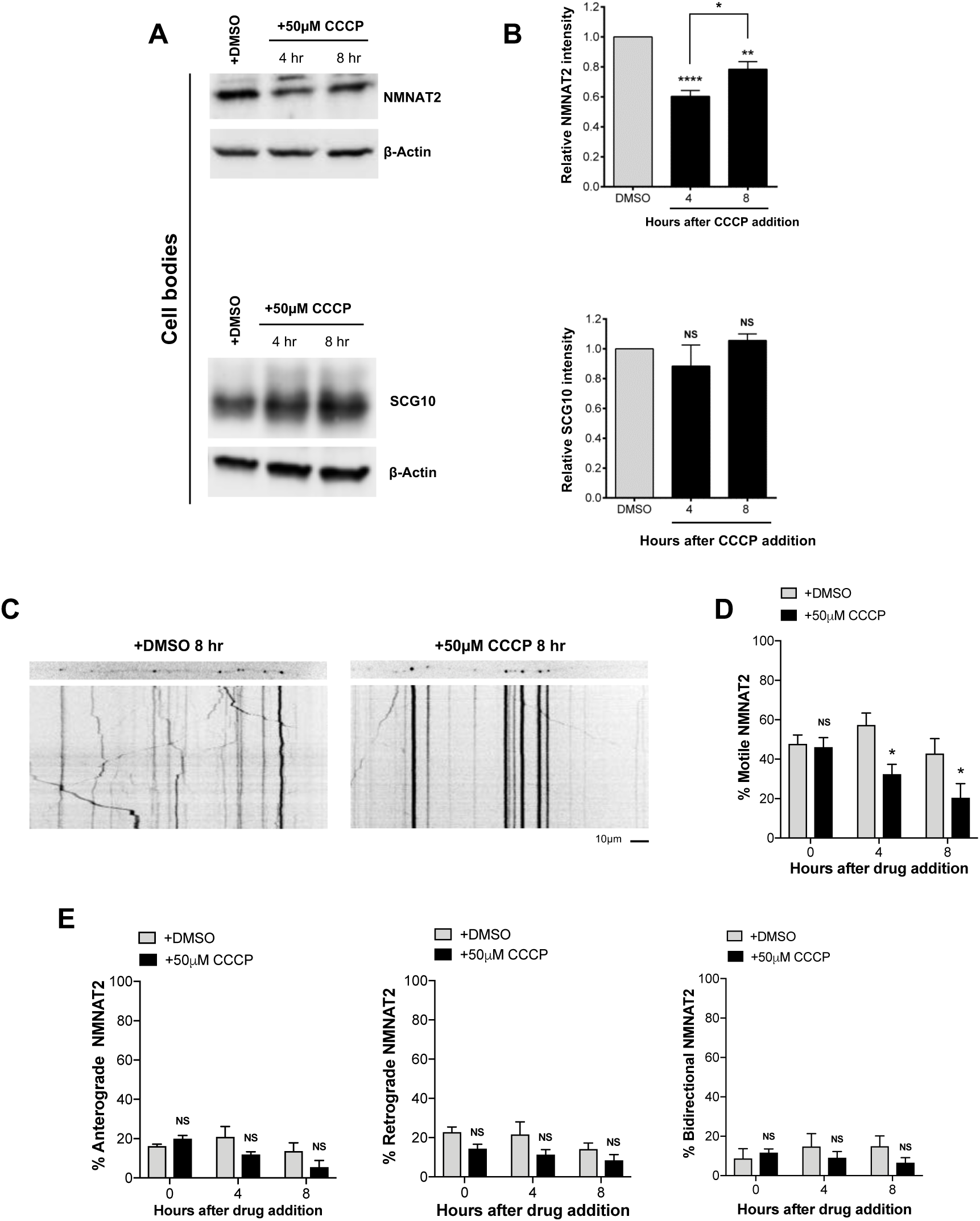
NMNAT2 depletion reflects impairment of both axonal transport and synthesis. **(A)** Representative immunoblot of wild-type SCG cell bodies/ganglia extracts probed for NMNAT2, SCG10 and β-actin (loading control) at the indicated time points after CCCP treatment. **(B)** Quantification of normalised NMNAT2 and SCG10 levels (to β-actin) is shown, with data presented relative to DMSO control (Mean±SEM; n=4; one-way ANOVA followed by Bonferroni post-hoc test; ****, p<0.0001; **, p<0.01; NS, non-significant). **(C)** Representative kymographs of wild-type SCG dissociated cultures expressing NMNAT2-EGFP. **(D)** Quantification of the % of motile NMNAT2 at the indicated time points after CCCP treatment from 3 neurites per condition in 4 independent experiments (Mean±SEM; n=4; two-way ANOVA followed by Sidak post-hoc test; *, p<0.05. NS, non-significant). **(E)** Quantification of the % of motile bidirectional, anterograde and retrograde NMNAT2 at the indicated time points after CCCP treatment from 3 neurites per condition in 4 independent experiments (Mean±SEM; n=4; two-way ANOVA followed by Sidak post-hoc test; NS, non-significant).

We next explored whether CCCP alters NMNAT2 axonal transport. We microinjected GFP-tagged NMNAT2 and followed changes in its axonal transport parameters. We found a significant reduction of the percentage of motile NMNAT2 vesicles at 4 and 8 hr after CCCP addition (Fig. 3C, D). This may also explain the slight recovery of NMNAT2 levels in cell bodies at 8 hr after CCCP addition following the decline at 4 hr (Fig. 3A, B), as any NMNAT2 that is synthesised would be less efficiently transported into neurites and would accumulate in cell bodies instead. The overall reduction in axonal transport of NMNAT2 appeared to be a result of a combination of impaired anterograde, retrograde and bidirectional transport, although separately none of the individual parameters reached statistical significance (Fig. 3E).

Thus, reduced axonal transport of NMNAT2 and reduced synthesis and/or enhanced degradation combine to reduce axonal NMNAT2 levels after CCCP treatment.

### Changes in the NMN/NAD ratio following CCCP administration

We have shown that NMNAT2 depletion leads to accumulation of its substrate, NMN, which we suggest promotes axon degeneration (Di Stefano et al., 2015, 2017; Loreto et al., 2015), as well as to NAD depletion, which also plays an important role (Essuman et al., 2017; Sasaki et al., 2016) (Fig. 4A). Thus, changes in NMN/NAD ratio is an additional indicator of Wallerian pathway activation. We previously reported a marked increase of NMN levels in injured sciatic nerves *in vivo* (Di Stefano et al., 2015, 2017). Sasaki and colleagues recently showed a transient increase in NMN levels in sensory neurons after axotomy also *in vitro* (Sasaki et al., 2016). However, selecting the correct time points is difficult due to the substantial cellular material required for the analysis and the rapid degeneration process which compromises the integrity of the plasma membrane, making any measurement unreliable. We therefore tested whether NMN accumulates and NAD declines following mitochondrial depolarisation in *Sarm1*^*-/-*^ SCG neurons, where the degeneration process following CCCP administration is strongly delayed (Fig. 1D, E). We looked at 12 hr after CCCP treatment, when wild-type neurites showed the first signs of degeneration (Fig. 2C, D), reasoning that an increase in NMN levels should have already occurred. We found a 2-fold increase in NMN levels and a more modest decrease in NAD levels in neurites resulting in a robust increase in the NMN/NAD ratio (Fig. 4B) (Fig. S1A), consistent with the predicted effects of NMNAT2 loss. In contrast, changes in the cell bodies were much more modest (Fig. 4C) (Fig. S1B), consistent with levels of NMNAT2 in the soma being less affected after CCCP administration (Fig. 3A, B) and with the presence of nuclear NMNAT1, which will contribute to NMN and NAD homeostasis in this compartment.

**Figure 4.**
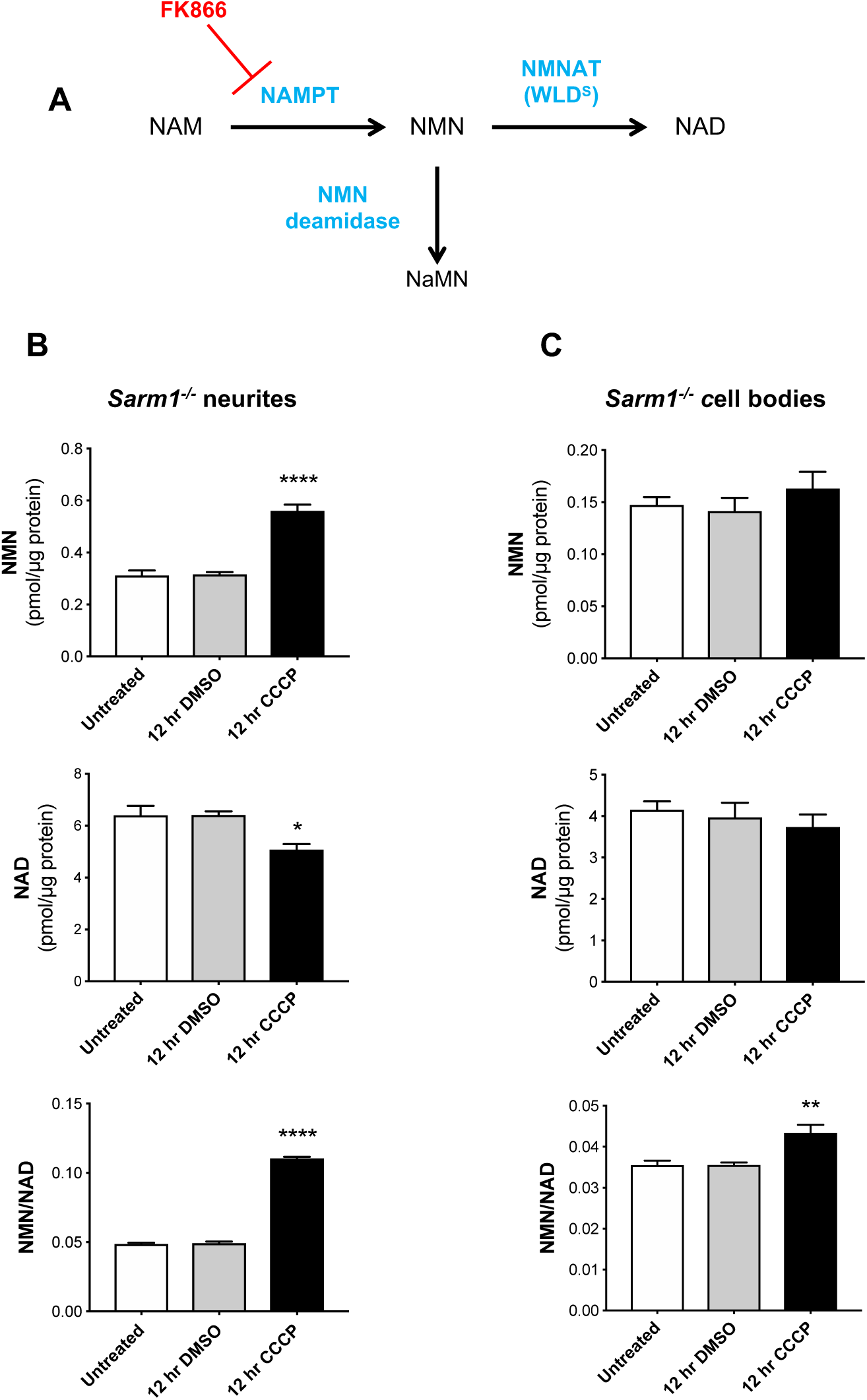
Changes in the NMN/NAD ratio following CCCP administration. **(A)** Schematic representation of NAD salvage pathway from nicotinamide and points at which FK866 and bacterial NMN deamidase will act (NAM, nicotinamide; NaMN, nicotinic acid mononucleotide; NMN, nicotinamide mononucleotide; NAD, nicotinamide adenine dinucleotide; NAMPT, nicotinamide phosphoribosyltransferase; NMNAT, nicotinamide mononucleotide adenylyltransferase). **(B, C)** NMN and NAD levels and NMN/NAD ratios in neurite (B) and cell body/ganglia (C) fractions from *Sarm1*^*-/-*^ SCG explant cultures at the indicated time points after CCCP treatment (Mean±SEM; n=5; one-way ANOVA followed by Bonferroni post-hoc test; ****, p<0.0001; **, p<0.01; *, p<0.05. Statistical significance shown relative to 12 hr DMSO).

Several lines of evidence suggest that NMN accumulation is not simply a marker but is a trigger of axon degeneration. Blocking NMN accumulation with FK866, an inhibitor of the NMN-synthesizing enzyme NAMPT (Fig. 4A), delays Wallerian degeneration. Exogenous administration of NMN restores its accumulation in the presence of FK866, reverting the protection (Di Stefano et al., 2015; Loreto et al., 2015). Also scavenging NMN with expression of bacterial enzyme NMN deamidase, which converts NMN into NaMN (Fig. 4A), results in strong protection of injured axons in mouse primary neurons and *in vivo* in mice and zebrafish (Di Stefano et al., 2015, 2017; Loreto et al., 2015). We therefore tested whether NMN accumulation also promotes axon degeneration after CCCP administration. We first confirmed that the levels of NAMPT were not affected by CCCP treatment (Fig. 5A, B). This is important since NAMPT expression is required for NMN synthesis, which results in the accumulation of NMN in the absence of NMNAT2. We then tested whether blocking NMN synthesis with FK866 delays CCCP-induced axon degeneration. As with axon degeneration after axotomy (Di Stefano et al., 2015), FK866 treatment strongly delayed neurite degeneration following CCCP administration. Of note, co-administration of exogenous NMN reverted FK866-induced protection (Fig. 5C, D). In contrast to our previous findings (Di Stefano et al., 2015), some studies reported a protective effect of NMN against axotomy-induced axon degeneration (Sasaki et al., 2006), possibly due to differences in incubation time of NMN before transection. Importantly, we confirmed that NMN had no protective effect on the degeneration process when added together with CCCP (Fig. 5E, F). NMNAT2 depletion still occurred in neurites protected by FK866, consistent with its expected protective action downstream of NMNAT2 loss in this situation (Fig. S2A) (Di Stefano et al., 2015). FK866 conferred full protection also when added up to 8 hr after CCCP addition (when NMNAT2 levels in neurites are already dramatically reduced) and halted the progression of the degeneration when added 12 hr after CCCP (when neurites appear already damaged) (Fig. S2B, C). This suggests that activation of the pathway might be reversible, or at least the existence of a time window after mitochondrial dysfunction when it can be prevented, which is important in the context of therapeutic intervention in human diseases.

**Figure 5.**
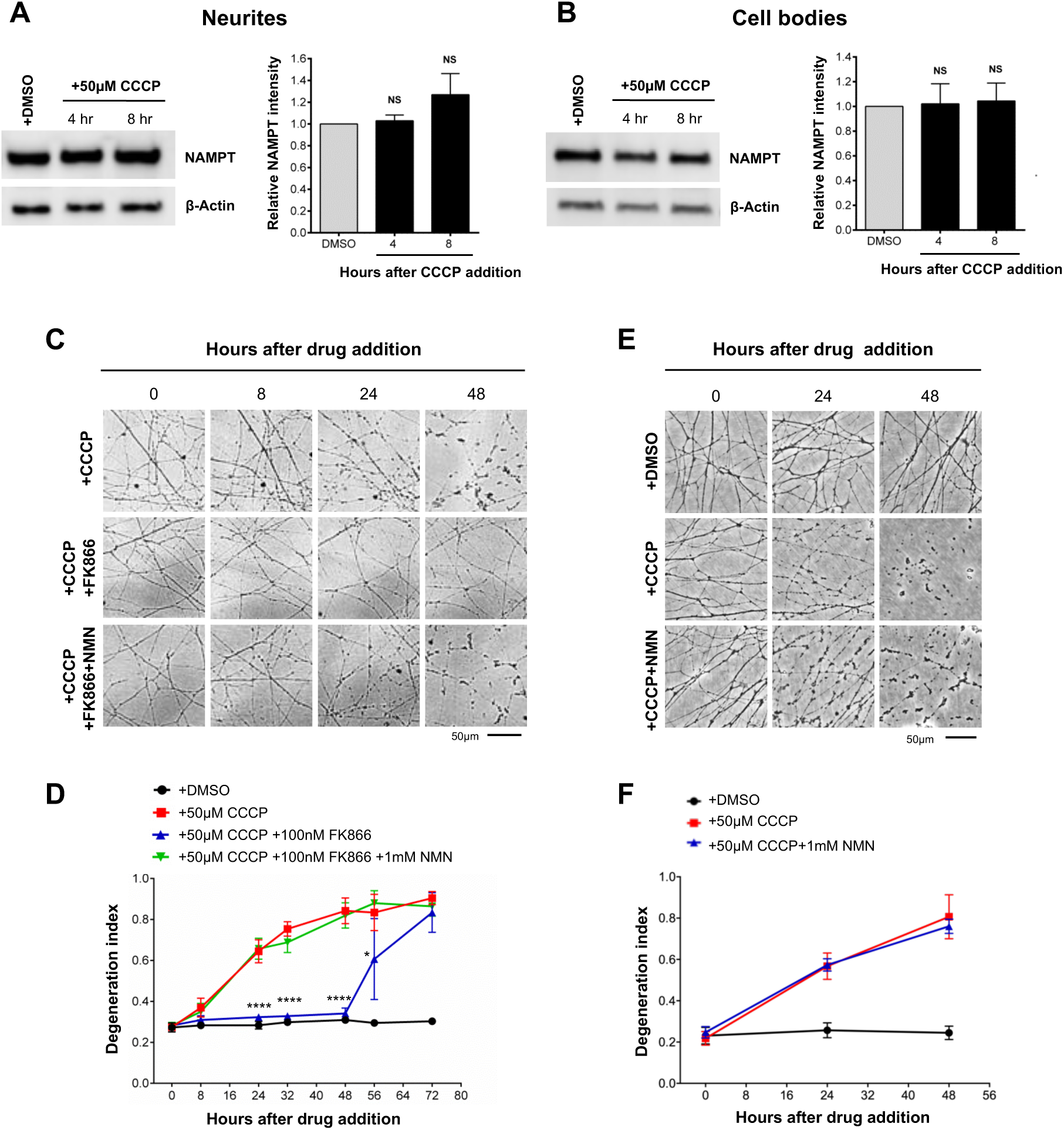
Inhibition of NMN synthesis protects neurites against CCCP toxicity. **(A, B)** Representative immunoblot of wild-type SCG neurite extracts probed for NAMPT and β-actin (loading control) at the indicated time points after CCCP treatment. Quantification of normalised NAMPT levels (to β-actin) is shown, with data presented relative to DMSO control (Mean±SEM; n=4; one-way ANOVA followed by Bonferroni post-hoc test; NS, non-significant). **(C)** Representative phase contrast images of neurites from wild-type SCG explant cultures at the indicated time points after CCCP, FK866 and NMN treatment. Where indicated, FK866 and NMN were added at the same time of CCCP. **(D)** Quantification of the degeneration index in experiments described in (C) from 3 fields per sample in 4 independent experiments (Mean±SEM; n=4; two-way ANOVA followed by Tukey post-hoc test; ****, p<0.0001. Statistical significance shown relative to +50µM CCCP). **(E)** Representative phase contrast images of neurites from wild-type SCG explant cultures at the indicated time points after CCCP and NMN treatment. **(F)** Quantification of the degeneration index in experiments described in (E) from 3 fields per sample in 4 independent experiments (Mean±SEM; n=4; two-way ANOVA followed by Tukey post-hoc test).

Taken together, these data further support a pro-degenerative role of NMN and are an additional confirmation that CCCP causes axon degeneration through the activation of the Wallerian pathway.

### *Highwire* deletion rescues loss of dopaminergic neurons in *Pink1 Drosophila* mutants

To validate our findings in an *in vivo* model where mitochondrial dysfunction is caused by a genetic mutation, we employed a *Drosophila* mutant with a loss-of-function mutation in the PD-associated gene *Pink1 (Pink1*^*B9*^). Pink1 is involved in mitochondrial quality control and mutations in this protein are linked to early-onset recessive PD (Pickrell and Youle, 2015; Valente et al., 2001, 2004). Loss of Pink1 in flies leads to severe mitochondrial defects resulting in, among other phenotypes, loss of dopaminergic neurons (in the PPL1 cluster), locomotor deficits and reduced lifespan (Clark et al., 2006; Hewitt and Whitworth, 2017; Park et al., 2006; Tain et al., 2009). The Wallerian pathway is evolutionary conserved, with several orthologous genes controlling axon degeneration both in mammals and flies (Freeman, 2014) (Fig. S3). As ubiquitous *dSarm* deletion is lethal in *Drosophila*, we instead opted to assess the effects of *Highwire* mutation on the *Pink1*^*B9*^ phenotype. Highwire, and its mammalian ortholog PHR1, are E3 ubiquitin ligases that target *Drosophila* NMNAT (dNMNAT) and NMNAT2, respectively, for proteasomal degradation and Highwire/PHR1 depletion appears to delay axon degeneration after axotomy by increasing levels and/or stabilising dNMNAT/NMNAT2, preventing the activation of the Wallerian pathway at an early step (Babetto et al., 2013; Xiong et al., 2012) (Fig. S3).

We first tested whether Highwire deficiency (*Hiw*^*ΔN*^) could rescue the loss of dopaminergic neurons in the PPL1 cluster (Fig. 6A) in *Pink1*^*B9*^ flies. As *Highwire* mutants display synaptic overgrowth during development at the neuromuscular junction (Wan et al., 2000), we first confirmed that the number of dopaminergic neurons in the PPL1 cluster did not differ from wild-type flies (Fig. 6B, C). Importantly, *Highwire* deletion rescued the loss of dopaminergic neurons in the PPL1 cluster (Fig. 6B, C). *Highwire* deletion also significantly prolonged the lifespan of *Pink1*^*B9*^ flies (Fig. 6D), but was not sufficient to rescue climbing and flying ability (Fig. 6E, F), likely due to the widespread muscle degeneration that is also seen in *Pink1*^*B9*^ flies (Clark et al., 2006; Tain et al., 2009). Modulation of the Wallerian pathway thus appears to be protective against neurodegeneration caused by non-toxin models of mitochondrial disruption in flies.

**Figure 6.**
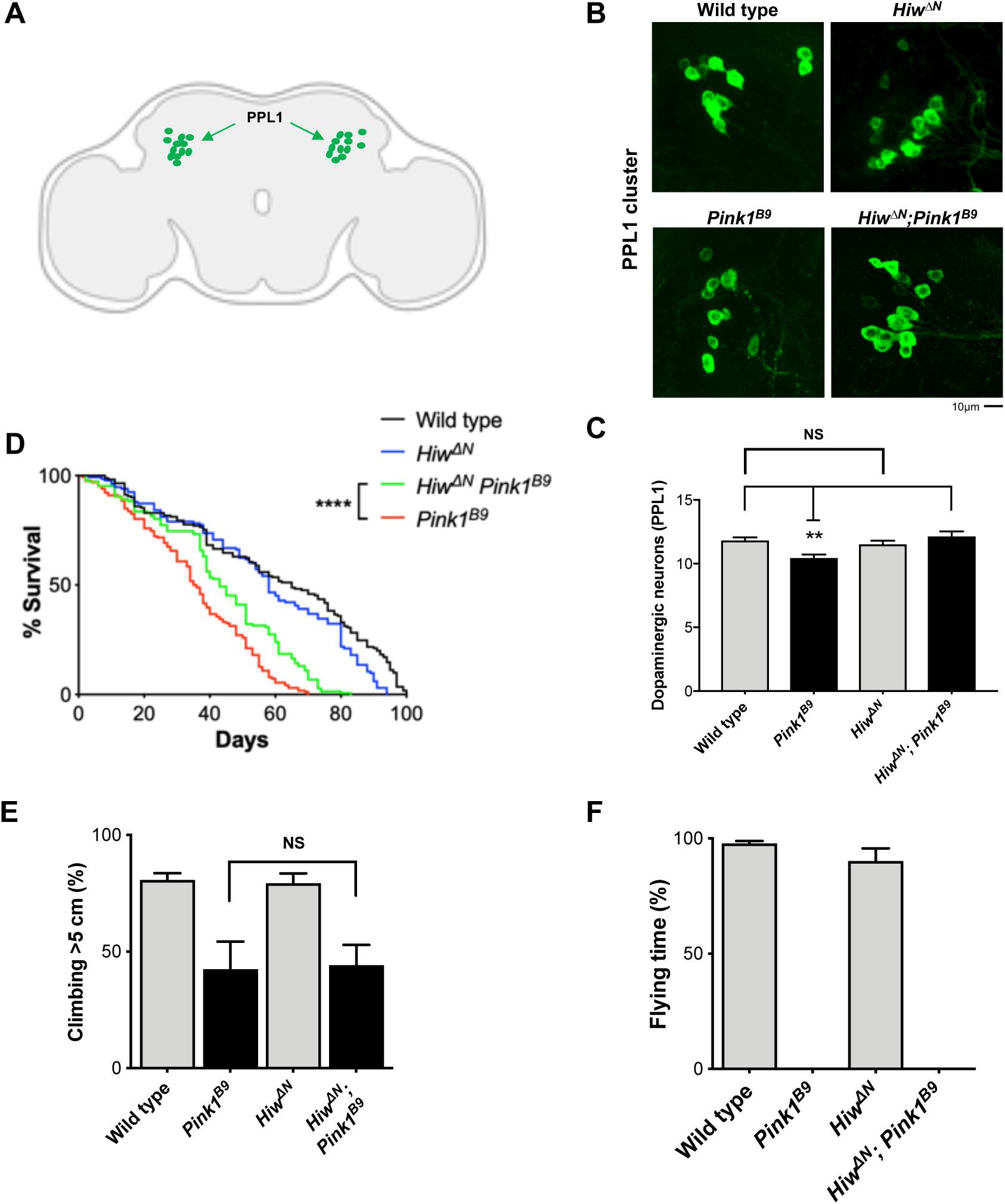
*Highwire* deletion rescues loss of dopaminergic neurons in *Pink1 Drosophila* mutants. **(A)** Schematic image of a *Drosophila* brain with the PPL1 cluster of dopaminergic neurons shown in green (‘Created with BioRender’). **(B)** Representative images of adult *Drosophila* (20 days old) brains stained with anti-TH antibody. The PPL1 cluster of dopaminergic neurons is shown. **(C)** Quantification of the number of dopaminergic neurons per PPL1 cluster (Mean±SEM; n=16-25; one-way ANOVA followed by Tukey post-hoc test; **, p<0.01). **(D)** Lifespan curves of wild-type, *Hiw*^*Δ*^*N, Pink1*^*B9*^, *Hiw*^*Δ*^*N Pink1*^*B9*^ flies (n>130 flies per condition; log-rank (Mantel-Cox) test. ****, p<0.0001). **(E, F)** Analysis of climbing and flying ability of 7 days old flies of the indicated genotypes (Mean±SEM; n=3 climbing, n=9 flying; one-way ANOVA followed by Tukey post-hoc test; NS, non-significant).

## DISCUSSION

The data presented here support an involvement of the Wallerian pathway in disorders involving mitochondrial dysfunction. First, acute mitochondrial depolarisation by CCCP leads to axon degeneration, in the absence of a physical injury, through the same pathway that regulates Wallerian degeneration. It does so by impairing axonal transport and synthesis (or stimulating degradation) of the axonal survival enzyme NMNAT2, leading to substantially reduced levels in neurites which increase the NMN/NAD ratio and trigger SARM1-dependent axon degeneration. In addition, neuroprotection of dopaminergic neurons conferred by *Highwire* deletion in flies carrying mutant Pink1 suggests a wider relevance of the Wallerian pathway to different types of mitochondrial insults *in vivo*.

Our previous work and that of others suggest a minor contribution of mitochondria to the late stages of Wallerian degeneration after axon transection (Kitay et al., 2013; Loreto et al., 2015), mainly through the opening of mitochondria permeability transition pore and release of Ca^2+^ into the cytoplasm (Barrientos et al., 2011; Villegas et al., 2014). We now show that mitochondrial dysfunction can impact on the Wallerian pathway in a second way, activating it at an early step upstream of NMNAT2. Crucially, like FK866-protected axons (Loreto et al., 2015), *Sarm1*^*-/-*^ and *Wld*^*S*^ axons can be kept morphologically intact for days despite fully depolarised mitochondria (this study and (Loreto et al., 2015; Summers et al., 2014)). This indicates that WLD^S^ expression and SARM1 deficiency confer protection downstream of mitochondrial impairment (Fig. 7), rather than directly impacting on mitochondrial health.

**Figure 7.**
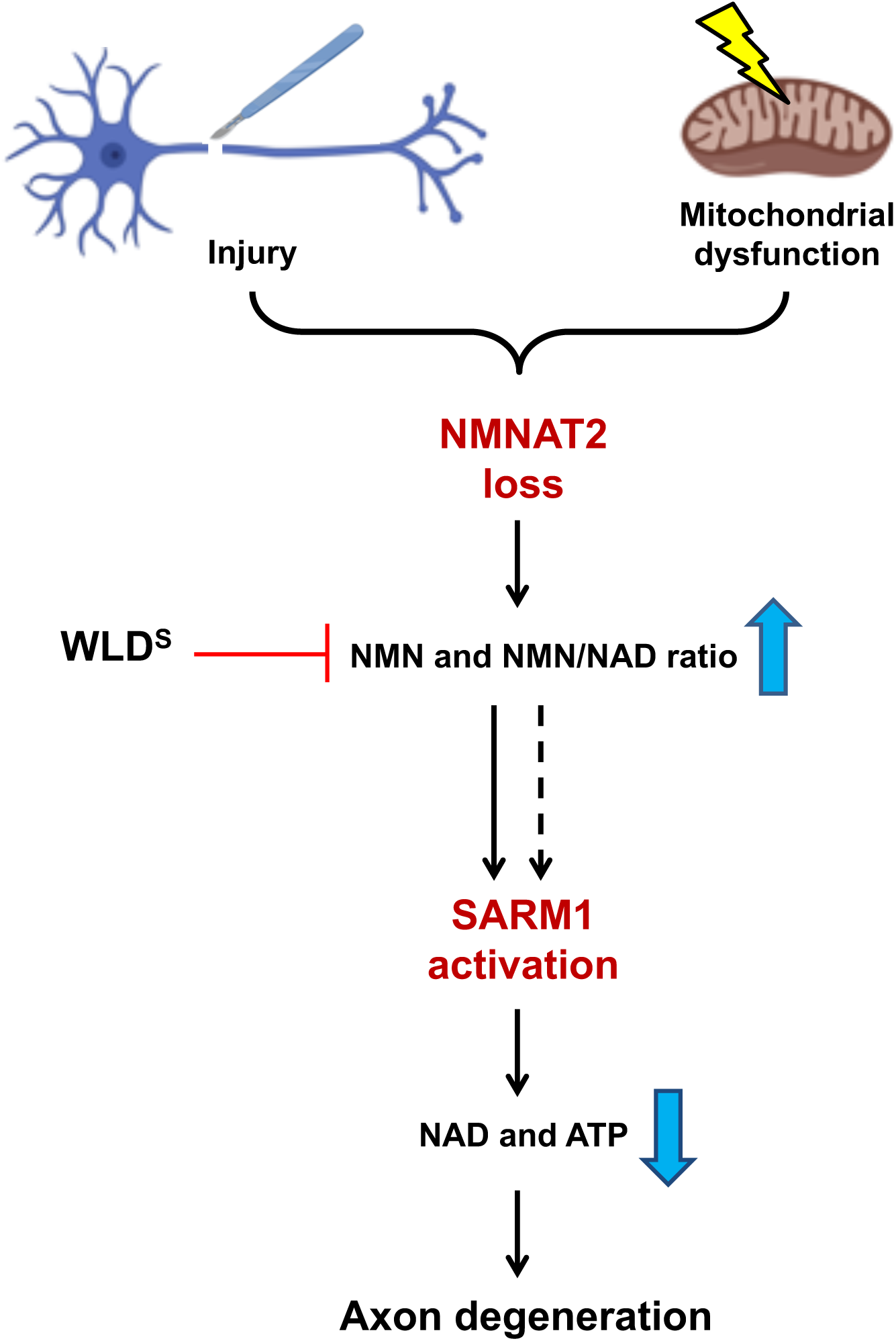
Mitochondrial dysfunction as an upstream signal activating the Wallerian pathway. Schematic representation of the Wallerian pathway (‘Created with BioRender’). Injury and mitochondrial impairment act as two independent insults resulting in the activation of the Wallerian pathway.

The relevance of the Wallerian pathway beyond its role after axotomy is now widely accepted and mitochondrial depolarisation can now be added to a growing list of non-axotomy insults causing Wallerian-like degeneration, including toxicity caused by chemotherapy agents, chemicals disruption of the nigrostriatal pathway, protein synthesis inhibition and NGF withdrawal (Conforti et al., 2014). Importantly, most of these studies used either WLD^S^ expression or *Sarm1* deletion as means to assess the involvement of Wallerian-like degeneration. However, these proteins are likely to have additional, non-Wallerian pathway functions and could thus confer a protective phenotype independently of the Wallerian pathway. For example, WLD^S^ protection against neuropathy and retinopathy in a streptozotocin-induced mouse model of diabetes is linked to a rescue of pancreatic islets (Zhu et al., 2011), likely through a mechanism that is unrelated to its role in axons. Recent steps forward in the understanding of the molecular mechanisms of axon degeneration revealed a well-defined pathway of axon death, with the identification of crucial mechanistic links between NMNAT2 and SARM1 (Gilley et al., 2015, 2017). The knowledge of a core mechanistic pathway allows multiple stages to be probed when seeking to establish a role for the Wallerian pathway in non-axotomy insults and diseases. Here, we followed this approach focusing on NMNAT2 levels, changes in NMN/NAD ratio and protection conferred by WLD^S^ expression and *Sarm1* deletion. This is the first demonstration of Wallerian pathway involvement at multiple steps in a non-axotomy axonal stress.

A next crucial question is whether the activation of the Wallerian pathway contributes to neurodegenerative disorders caused by mitochondrial dysfunction. CCCP is widely used to impair mitochondrial function and has proven instrumental for understanding the role of mitochondria in a number of physiological and non-physiological cellular processes. However, it remains unclear how much its potent and acute mitochondrial toxicity reflects chronic mitochondrial dysfunction in human pathologies. The strong protection achieved by blocking the Wallerian pathway is remarkable, but the extent of mitochondrial damage in neurodegenerative disorders is likely to be milder. The neuroprotection *in vivo* in *Pink1* mutant flies represents a first indication of the possible wider relevance of the Wallerian pathway to other mitochondrial insults *in vivo*, although the use of alternative means to impair mitochondria could provide further understanding of the mechanisms involved. The protection of neuronal soma in *Pink1*^*B9*^ flies could be secondary to rescue of axon loss. Conversely, *Drosophila* only has one NMNAT isoform (compared to three in mammals) and so a reduction in dNMNAT levels would likely cause a more profound damage to the whole cell, rather than predominantly affecting axons (as it is the case with the major axonal isoform, NMNAT2, in mammals). Finally, we cannot fully rule out the possibility that other actions of Highwire contribute to these observations.

Among a number of neurodegenerative disorders associated with mitochondrial dysfunction, the link between PD and axon loss is particularly important. PD involves preferential loss of substantia nigra pars compacta dopaminergic neurons. These neurons have extremely long and branched axons which are lost early in PD patients (Matsuda et al., 2009; Tagliaferro and Burke, 2016), and, as such, may be more vulnerable to axonal stresses. Wallerian-like degeneration has also been implicated in other PD models, with WLD^S^ protecting after MPTP and 6-hydroxydopamine administration (Cheng and Burke, 2010; Hasbani and O’Malley, 2006; Sajadi et al., 2004), and with neuroprotection in *Pink1* mutant flies by Highwire deficiency that can now be added to the list. However, more comprehensive studies in genetic and chronic models of PD in mammals will be needed to establish whether the Wallerian pathway plays a causative role in PD pathology or simply increases susceptibility to disease. Interestingly, we also show that lower levels of NMNAT2 make neurites more vulnerable to the consequences of CCCP-induced mitochondrial depolarisation and, as NMNAT2 mRNA levels have been reported to vary hugely in the human population (up to 50-fold differences) (Ali et al., 2016), some individuals might thus be at a much higher risk of mitochondrial disorders.

To conclude, we show that acute mitochondrial impairment induced by CCCP leads to NMNAT2 depletion and subsequent activation of the Wallerian pathway (Fig. 7), and that loss of dopaminergic neurons as a result of mitochondrial dysfunction in flies with *Pink1* loss-of-function mutation can be prevented by modulation of the Wallerian pathway by *Highwire* deletion. This study provides mechanistic insights on how mitochondrial dysfunction leads to axon degeneration and identifies the Wallerian pathway as a potential contributor to axon pathology in mitochondrial disorders. It is now important to test the role of the pathway in models that more closely replicate human mitochondrial diseases.

## MATERIALS AND METHODS

All studies conformed to the institution’s ethical requirements in accordance with the 1986 Animals (Scientific Procedures) Act.

### Primary neuronal cultures

C57BL/6J or CD1 (referred to as wild-type, Charles River, UK), *Wld*^*S*^, *Nmnat2*^*+/+*^, *Nmnat2*^*+/gtE*^, *Nmnat2*^*gtBay/gtE*^ and *Sarm1*^−/−^ mouse SCG explants were dissected from P0-2 pups. Explants were cultured in 35 mm tissue culture dishes pre-coated with poly-L-lysine (20 µg/ml for 1 hr; Sigma) and laminin (20 µg/ml for 1 hr; Sigma) in Dulbecco’s Modified Eagle’s Medium (DMEM, Gibco) with 1% penicillin/streptomycin, 100 ng/ml 7S or 50 ng/ml 2.5S NGF (all Invitrogen) and 2% B27 (Gibco). 4 µM aphidicolin (Merck) was used to reduce proliferation and viability of small numbers of non-neuronal cells. For cultures of dissociated SCG neurons, wild-type SCG explants were incubated in 0.025% trypsin (Sigma) in PBS (without CaCl_2_ and MgCl_2_) (Sigma) for 30 min followed by incubation with 0.2% collagenase type II (Gibco) in PBS for 20 min. Ganglia were then gently dissociated using a pipette. Dissociated neurons were plated in a poly-L-lysine and laminin-coated area of ibidi μ-dishes (Thistle Scientific) for microinjection experiments. Dissociated cultures were maintained as explant cultures except that B27 was replaced with 10% fetal bovine serum (Sigma). Culture media was replenished every 3 days. Neurites were allowed to extend for 7 days before performing the experiments.

### Drug treatments

Uncut SCG neurons were treated with CCCP or vehicle (DMSO) just prior to imaging (time 0 hr). Unless specified, FK866 (kind gift of Prof. Armando Genazzani, University of Novara) and NMN (Sigma) were added at the same time as CCCP. The incubation time and the drug concentration used for every experiment are indicated in the figures and/or figure legends.

### Acquisition of phase contrast images and quantification of axon degeneration

Phase contrast images were acquired on a DMi8 upright fluorescence microscope (Leica microsystems) coupled to a monochrome digital camera (Hammamatsu C4742-95) or on a Zeiss TIRF microscope coupled to an EMCCD (Photometrics PVCam) camera using Axiovision software (Carl Zeiss Inc.). The objectives used were HCXPL 20X/0.40 Corr and Zeiss EC Plan Neofluar 20X/0.5 NA. The axon degeneration index (Sasaki et al., 2009) was determined using an ImageJ plugin (Schneider et al., 2012) (http://rsb.info.nih.gov/ij/download.html) which calculates the ratio of fragmented axon area over total axon area after binarization of the pictures and subtraction of the background.

### Determination of ATP levels

For measurement of ATP levels, dissociated SCG neurons were plated in 96-well plates at the same density. ATP measurements were performed with the ATPlite Luminescence Assay System (PerkinElmer). Two technical repeats were performed per each condition for every experiment. Data are expressed as % relative to DMSO control.

### Western blot

Following treatment with CCCP, SCG ganglia were separated from their neurites with a scalpel. Neurites originating from 15 ganglia were collected per condition, washed in ice-cold PBS containing protease inhibitors (Sigma), and lysed directly in 15 µl 2x Laemmli buffer containing 10% 2-Mercaptoethanol (Sigma). The remaining 15 ganglia were also collected and lysed. For NMNAT2 immunoblots, 14 µl of protein samples were loaded on a 12% SDS polyacrylamide gel. For SCG10 and NAMPT immunoblots, 1:15 dilutions of the original samples were loaded on a 12% SDS polyacrylamide gel. Membranes were blocked for 3 hr in 5% milk in TBS (50 mM Trizma base and 150 mM NaCl, PH 8.3, both Sigma) plus 0.05% Tween-20 (Sigma) (TBST), incubated overnight with primary antibody in 5% milk in TBST at 4°C and subsequently washed in TBST and incubated for 1 hr at room temperature with HRP-linked secondary antibody (Bio-Rad) in 5% milk in TBST. Membranes were washed, treated with ECL (Enhanced Chemiluminescence detection kit; Thermofisher) and imaged with Uvitec Alliance imaging system. The following primary antibodies were used: mouse anti-NMNAT2 (WH0023057M1 Sigma, 2 µg/ml), mouse anti-NAMPT (clone OMNI 379, Cayman Chemical Company, 1:2000) and rabbit anti-SCG10 (10586-1-AP Proteintech, 1:3000). Mouse anti β-actin was used as a loading control (A5316 Sigma, 1:5000). Quantification of band intensity was determined by densitometry using ImageJ.

### NMNAT2 axonal transport

Dissociated SCG neurons were microinjected using a Zeiss Axiovert S100 microscope with an Eppendorf FemtoJet microinjector and Eppendorf TransferMan^®^ micromanipulator. Plasmids were diluted in 0.5x PBS (without CaCl_2_ and MgCl_2_) and filtered using a Spin-X filter (Costar). The mix was injected directly into the nuclei of SCG neurons using Eppendorf Femtotips. Approximately 100 neurons were injected per dish. Injected plasmids were allowed to express for 16 hr before CCCP treatment. Plasmids were injected at the following concentrations: 30 ng/µl NMNAT2-EGFP, 30 ng/µl pDsRed2-N1. Time-lapse imaging of axonal transport was performed on an Olympus IX70 imaging system with 100X/1.35 Oil objective. During imaging, cell cultures were maintained at 37°C and 5% CO_2_ in an environment chamber. Images were captured at 4 frames per sec for 2 min. Three neurites per condition were imaged in every individual experiment.

### Determination of NMN and NAD levels

Following treatment with CCCP, *Sarm1*^*-/-*^ SCG ganglia were separated from their neurites with a scalpel. Neurites and cell bodies were washed in ice-cold PBS and rapidly frozen in dry ice and stored at −80 °C until processed for measuring NMN and NAD. Briefly, pyridine and adenine nucleotides were extracted by sonication in HClO_4_ in the presence of cAMP (as internal standard) and subsequently analysed by ion pair C18-HPLC chromatography and by spectrofluorometric HPLC analysis after derivatization with acetophenone (Mori et al., 2014). The levels of NMN and NAD were normalised to protein levels.

### *Drosophila* experiments

Newly enclosed flies were collected daily and separated by sex into vials of 20-35 flies for aging and experimental use. Genotypes used are *w*^*1118*^ (wild-type), *Pink1*^*B9*^, *Hiw*^*ΔN*^ and *Hiw*^*ΔN*^ *Pink1*^*B9*^. All flies were maintained at a constant 25°C temperature and humidity, in plastic vials with standard agar/cornmeal/yeast feed. Flies were exposed to a 12 hr light-dark cycle. All experiments were conducted on male flies. For PPL1 dopaminergic neuron staining, fly brains were dissected in cold 1x PBS and fixed in 4% paraformaldehyde-PBS (Sigma) for 30 min. Samples were washed in 1x PBS with 0.3% Triton X-100 (Sigma) and blocked for 1 hr at room temperature in 1x PBS with 0.3% Trition X-100 and 1% BSA (Sigma). Brains were incubated in primary antibody for 72 hr. After washing and incubation in a fluorescent secondary antibody solution for 4 hr, samples were mounted between two coverslips in ProLong diamond antifade mountant (ThermoFisher). Confocal images were acquired on a Leica microscopy system and blinded for analysis. Antibodies used were mouse anti-Tyrosine Hydroxylase 1:100 (22941, Immunostar Inc.) and secondary anti-mouse IgG (H+L) Alexa Fluor 488 (A11034, ThermoFisher). Flight assay was performed as previously described (Agrawal and Hasan, 2015). Briefly, flies were anaesthetised on ice for 5 min; the flat of a 30G 1” needle (Sigma) was attached to the anterior notum of a fly just posterior to the neck using clear nail varnish, leaving flight muscles unimpeded. Flies were given 15 min to recover. Needles were fixed in place under a video microscope. If required, a gentle mouth-blown puff of air was used to stimulate flight and the flying time was recorded for 30 sec. This was repeated 3 times per fly and the average of time spent in flight was calculated for each condition. For climbing assays, flies were gently transferred to fresh empty polystyrene vials without anaesthesia with a maximum density of 25 flies per vial. Groups of up to 6 vials were inserted into the RING device and after 5 min for the flies to adjust to the environmental change the device was tapped three times to settle flies to the bottom of the vials. 5 sec after the last tap, a picture was taken to assess the height climbed. Maximum height achieved was graded into 5 mm intervals, flies that climbed less than 5 cm were scored zero, and any fly that exceeded 5 cm was awarded the maximum score. This was repeated 3 times at 60 sec intervals and an average score given for that vial.

### Statistical analysis

Appropriate statistical testing of data was performed using Prism (GraphPad Software, La Jolla, USA). ANOVA with Tukey’s, Sidak’s or Bonferroni’s post hoc correction (as applicable), and log-rank (Mantel-Cox) test were used in this study. The n numbers in each individual experiment and the tests used are described in the figure legends. A p value < 0.05 was considered significant (****, p<0.0001; ***, p<0.001; **, p<0.01; *, p<0.05; NS, non-significant).

## AUTHOR CONTRIBUTIONS

A.L., L.C. and M.P.C conceived the study. A.L. designed and conducted most experiments and data analysis. C.S.H., V.L.H., A.S-M., and A.J.W. performed experiments on flies. G.O. and C.A. performed nucleotide measurements and related data analysis. C.L. helped with western blots. F.D.-B and M.P.C supervised and co-ordinated the research. A.L., F.D.-B and M.P.C. wrote the manuscript, with input from J.G..

## ACKNOWLEDGMENTS

We thank the members of the Coleman, Conforti and Dajas-Bailador lab for useful discussion. We thank Dr Jemeen Sreedharan for advice on fly experiments. This work was funded by the Faculty of Medicine and Health Sciences, School of Life Sciences (University of Nottingham), a Parkinson’s UK grant [grant number G-1602], the UK Medical Research Council [grant number MR/N004582/1 and MC_UU_00015/6], a Wellcome Trust PhD Fellowship for Clinicians and a Sir Henry Wellcome postdoctoral fellowship from the Wellcome Trust [grant number 210904/Z/18/Z].

