## Supplementary material for "Mitochondrial impairment activates the Wallerian pathway through depletion of NMNAT2 leading to SARM1-dependent axon degeneration": Fig. S1, Fig. S2, Fig. S3

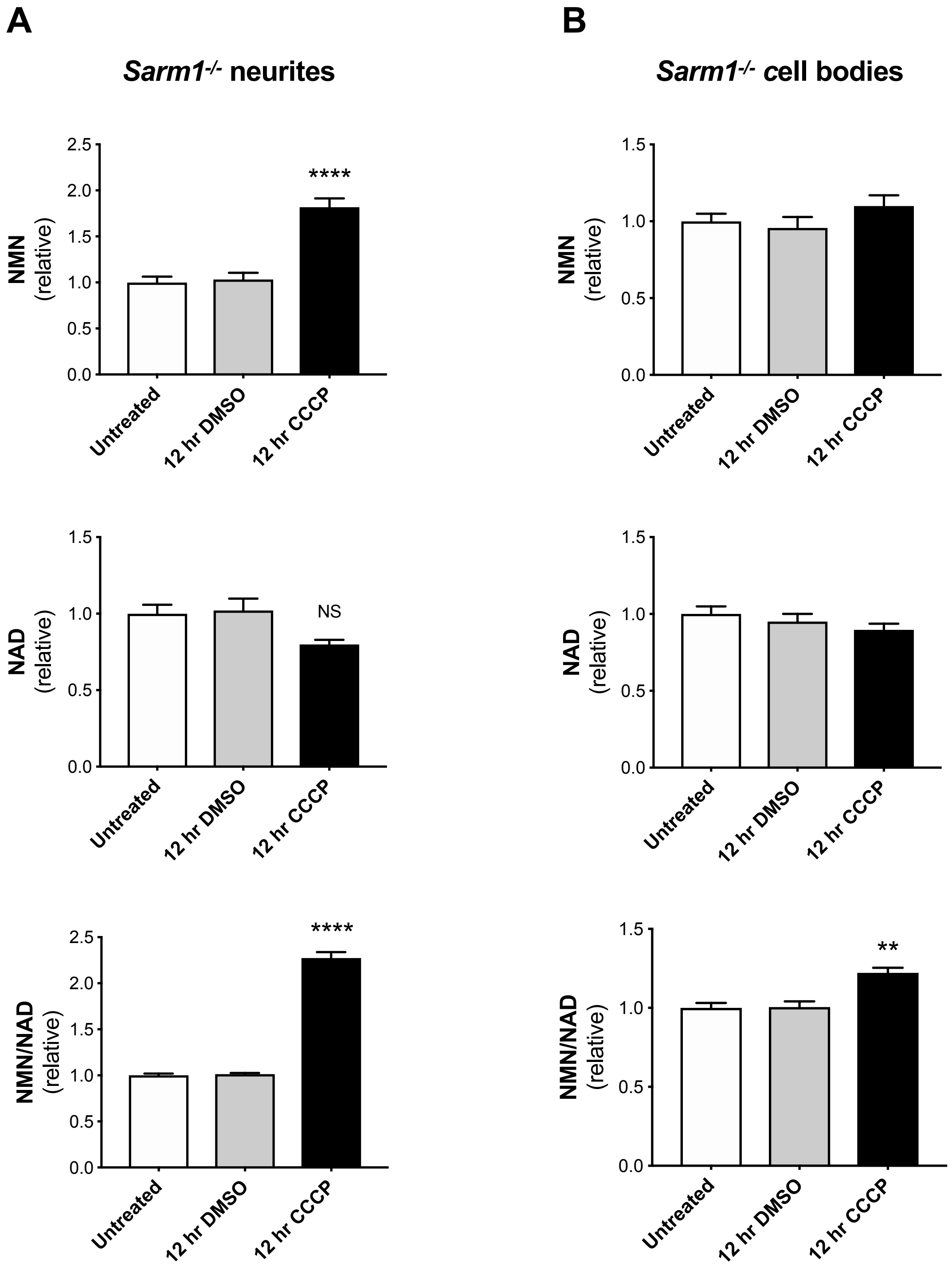


**Figure S1. (A,B)** Relative changes in NMN, NAD and NMN/NAD ratios in separate neurite (A) and cell bodies/ganglia (B) fractions from *Sarm1*$^{-/-}$ SCG explant cultures at the indicated time points after CCCP treatment (Mean±SEM; n=5; one-way ANOVA followed by Bonferroni post-hoc test; ****, $p<0.0001$; **, $p<0.01$; NS, non-significant. Statistical significance shown relative to 12 hr DMSO).

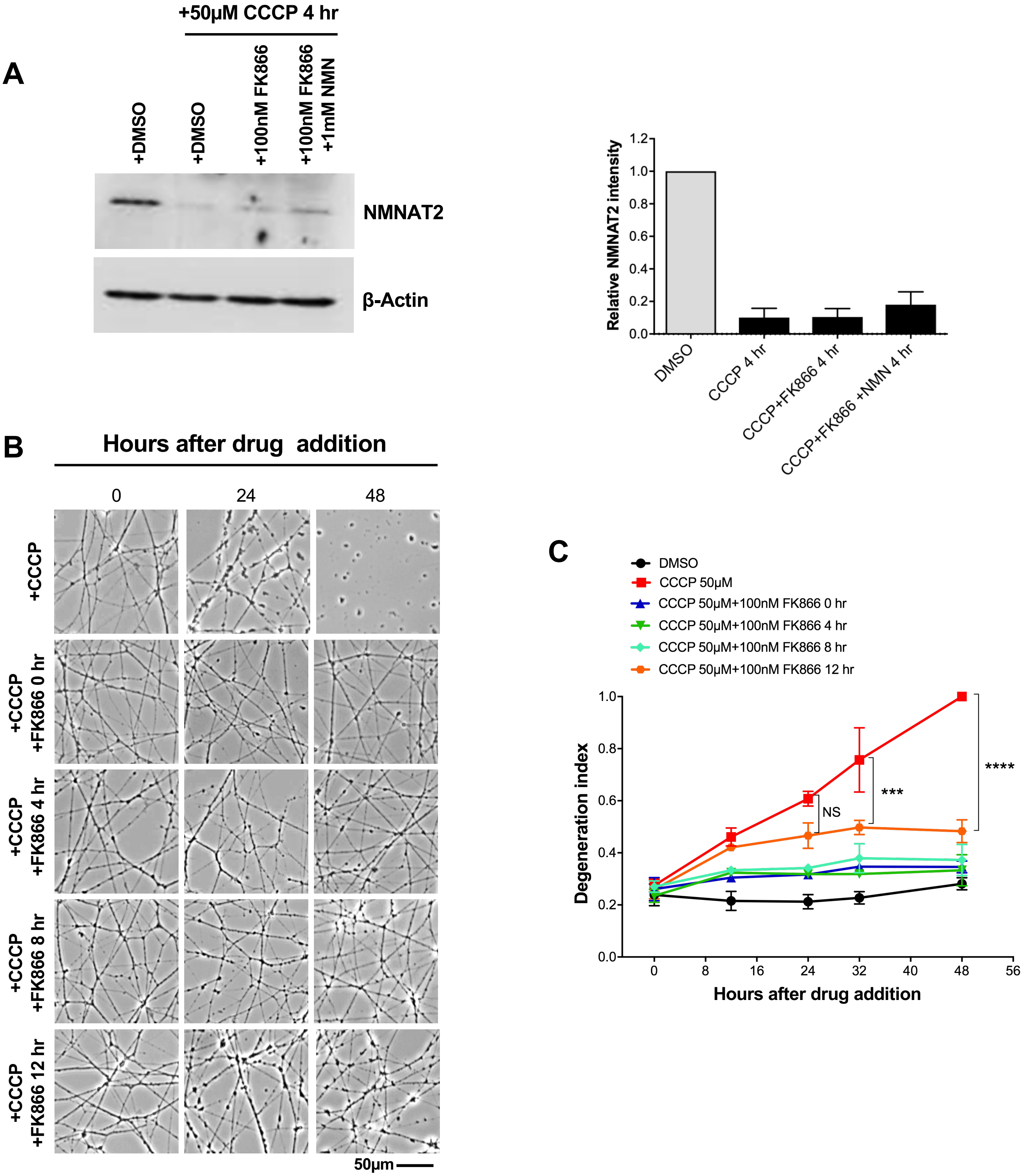


**Figure S2. (A)** Representative immunoblot of wild-type SCG neurite extracts probed for NMNAT2 and β-actin (loading control) 4 hr after CCCP, FK866 and NMN treatment. Quantification of normalised NMNAT2 and SCG10 levels (to β-actin) is shown, with data presented relative to DMSO control (Mean±SEM; n=2). **(B)** Representative phase contrast images of neurites from wild-type SCG explant cultures at the indicated time points after CCCP treatment. Where indicated, FK866 was added either at the same time of CCCP (0 hr) or 4, 8, 12 hr after CCCP addition. **(C)** Quantification of the degeneration index in experiments described in (B) from 3 fields per sample in 3 independent experiments (Mean±SEM; n=3; two-way ANOVA followed by Tukey post-hoc test; ****, $p<0.0001$; ***, $p<0.001$; NS, non-significant. Statistical significance shown relative to +50µM CCCP).

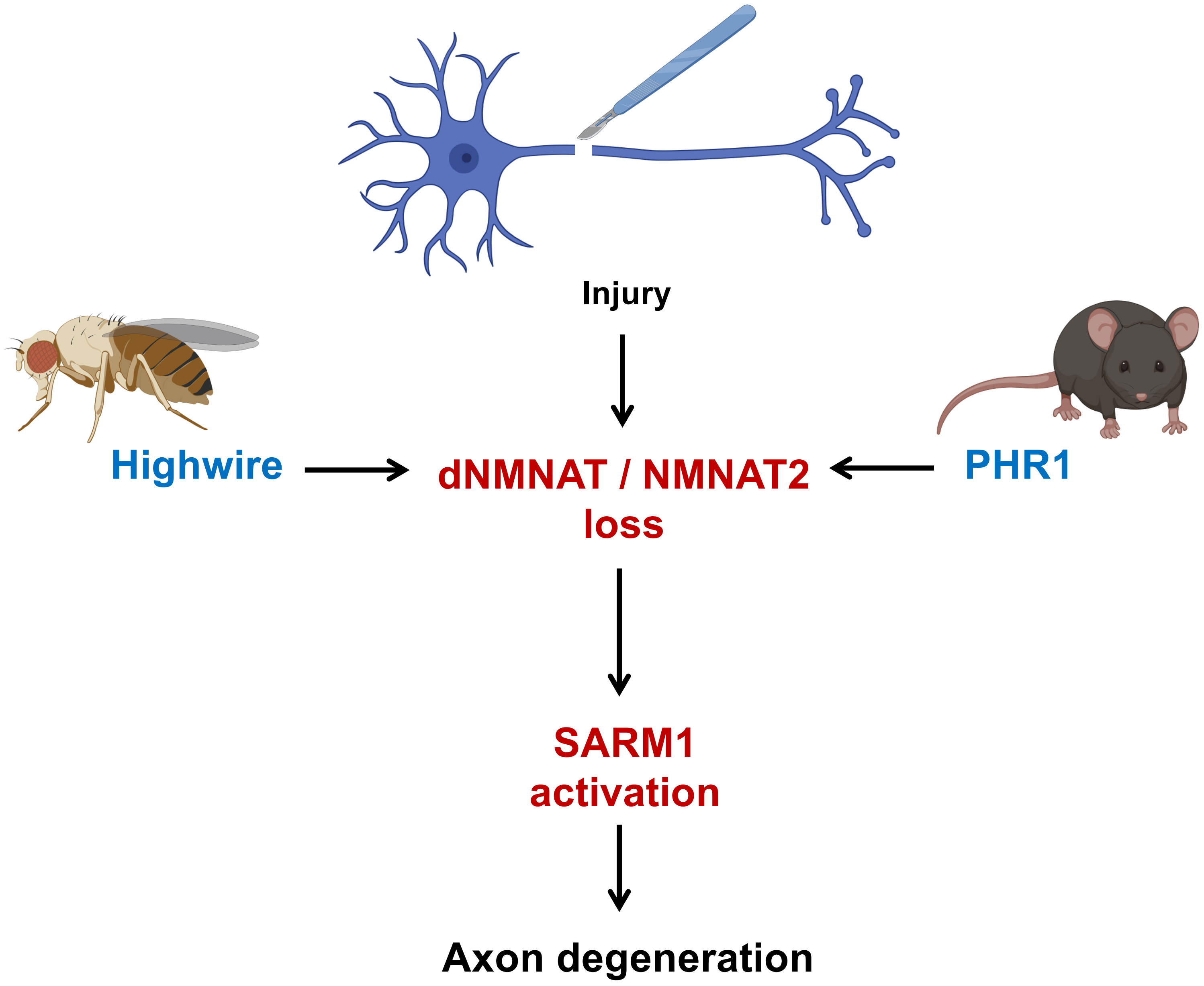


**Figure S3.** The Wallerian pathway is evolutionary conserved in flies and mammals. Highwire position in the pathway is shown ('Created with BioRender').
